# Genomic responses to selection for tame/aggressive behaviors in the silver fox (*Vulpes vulpes*)

**DOI:** 10.1101/228544

**Authors:** Xu Wang, Lenore Pipes, Lyudmila N. Trut, Yury Herbeck, Anastasiya V. Vladimirova, Rimma G. Gulevich, Anastasiya V. Kharlamova, Jennifer L. Johnson, Gregory M. Acland, Anna V. Kukekova, Andrew G. Clark

## Abstract

Animal domestications have led to a shared spectrum of striking behavioral and morphological changes. To recapitulate this process, silver foxes have been selectively bred for tame and aggressive behaviors for over 50 generations at the Institute for Cytology and Genetics in Novosibirsk, Russia. To understand the genetic basis and molecular mechanisms underlying the phenotypic changes, we profiled gene expression level and coding SNP allele frequencies in two brain tissues from 12 aggressive and 12 tame foxes. Expression analysis revealed 146 genes in prefrontal cortex and 33 genes in basal forebrain that were differentially expressed (5% FDR). These candidates include genes in key pathways known to be critical to neurological processing, including the serotonin and glutamate receptor pathways. In addition, 295 of the 31,000 exonic SNPs show significant allele frequency differences between tame and aggressive population (1% FDR), including genes with a role in neural crest cell fate determination.

## Introduction

Differences in the behavior of domesticated animals from their wild ancestors provide some of the best examples of the influence of genes on behavior (1). Domesticated animals have been selected to be easy to handle, and they generally exhibit reduced aggressiveness and increased social tolerance to both humans and members of their own species (2). Even after genomes of most domesticated species and their wild ancestral species have been sequenced, the identification of genes responsible for these behavioral differences has proven to be challenging (3-6). The selection for different traits in each of the domesticated animals and the antiquity of the time frame make it difficult to identify which genetic changes are causally responsible for changes in behavior (3, 7, 8).

Unlike the species domesticated historically, the silver fox (a coat color variant of the red fox, *Vulpes vulpes*) has been domesticated under controlled farm conditions at the Institute of Cytology and Genetics (ICG) of the Russian Academy of Sciences (9-11).The red fox and the domestic dog (*Canis familiaris*) share a common ancestor just 10 million years ago (12), making the fox experiment a model for dog domestication. To test whether selection for behavior was the primary force in the canine domestication process, starting in 1959, Drs. Dmitry Belyaev and Lyudmila Trut have been selecting conventional farm-bred foxes against fear and aggression to humans, followed by selection for contact-seeking behavior, which led to the development of a tame strain of foxes (Figure 1A) (9-11). The response to selection was extremely rapid: the first tame animal classified as “elite of domestication” appeared in generation 4, 1.8% of such foxes were observed at generation 6 (4/213), and by generation 45 almost all foxes belonged to that category (11). Foxes from the tame population relate with humans in a positive manner similar to that of friendly dogs (13). They are eager to establish human contact by one month after birth, and remain friendly throughout their entire lives (11).

**Figure 1.**
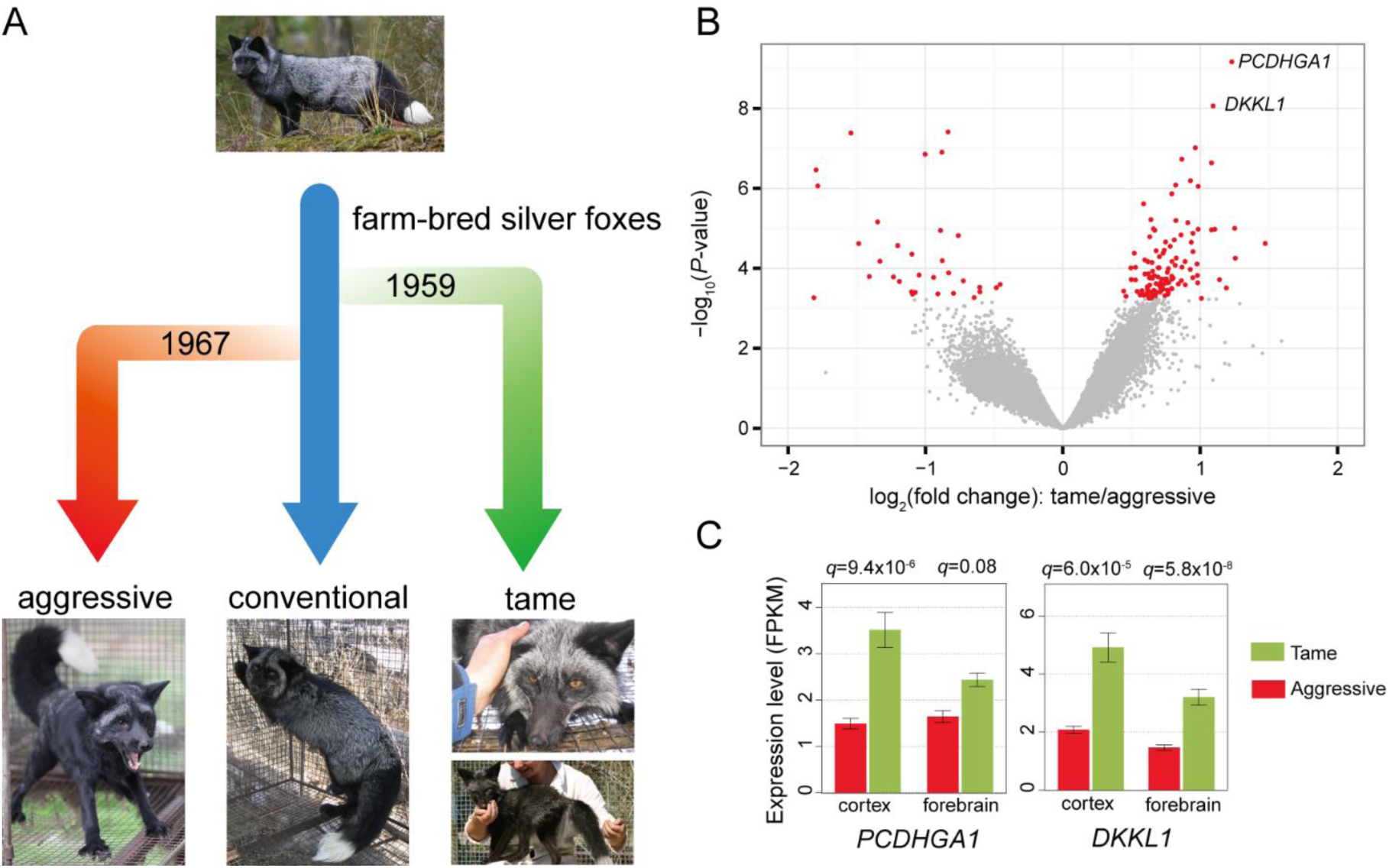
RNA-seq analysis identified differentially expressed genes in brain tissues between tame and aggressive fox population. (**A**) Artificial selection scheme for tameness and aggression in foxes. The conventional population of farm-bred foxes (blue arrow) was a founding population for both tame and aggressive fox populations. The population of conventional farm-bred foxes is still maintained in Novosibirsk. Since 1959, the selection experiment for tame foxes has been carried out to recreate the evolution of canine domestication. In 1970, an aggressive population was also selected to compare with the tame population. (**B**) A volcano plot showing differentially expressed genes detected in 12 tame and 12 aggressive fox prefrontal cortex samples. Plotted on the *x*-axis is the log_2_ fold difference between tame and aggressive samples. Plotted on the *y*-axis is –log_10_(*P*-value) calculated with the R package edgeR. Significant differentially expressed genes (FDR<0.05) are indicated in red and non-significant genes in gray. (**C**) Barplot of RNA-seq expression level with *q*-value in prefrontal cortex and forebrain samples for the top two significant candidate genes: *PCDHGA1* and *DKKL1*.

In parallel with selection for tameness, selective breeding for aggressive response to humans was started in 1970, with the aim to develop a population demonstrating less variation in behavior than conventional foxes (10, 11). This trait also showed a selection response (Figure 1A). The tame and aggressive fox strains were selected solely for specific behavioral traits, and the pedigree information was maintained during the entire breeding program (10, 11). Efforts were made to avoid close inbreeding in these populations, allowing continuous selection for many decades and generations (9-11). The heritability of these behavioral traits has been confirmed in multiple experiments (14-17), making these fox strains a promising model for the identification of the genetic basis of tame and aggressive behaviors.

To identify the genetic basis of the behavioral differences between tame and aggressive fox strains we developed the fox meiotic linkage map, experimental cross-bred pedigrees, and mapped eight significant and suggestive quantitative trait loci (QTL) for behavioral traits (17-20). Although QTL mapping is a promising strategy for the identification of genomic regions implicated in complex traits, this approach alone usually does not allow identification of the causative genes and mutations. In the current study we analyzed fox brain transcriptomes of 12 aggressive and 12 tame individuals. We evaluated gene expression in two brain regions: prefrontal cortex and basal forebrain. Prefrontal cortex is the site of memory and learning. It coordinates a wide range of neural processes and plays a central role in the synthesis of diverse information needed for complex behavior (21). The tamable animals may have altered learning abilities due to gene expression changes in the prefrontal cortex. Basal forebrain modulates cortical activity and plays an important role in arousal, attention, decision-making (22). RNA-seq analysis of these two brain regions identified significant differences in gene expression between the two fox strains and pinpointed several gene networks that were modified in the course of artificial selection for tame/aggressive behaviors.

## Results and Discussion

### Gene expression profile in the brain altered after selection for tameness

The profound behavior differences happened rapidly after selection, and brain gene expression level changes might play an important role in the response. To investigate this, Illumina RNA-seq experiments were performed on brain tissue from 12 aggressive and 12 tame individuals (Figures S1-S3), including the right prefrontal cortex and right basal forebrain (Figure S4). These experiments yielded a total of 1.57 billion RNA-seq reads, with an average of 30 million reads per sample (Table S1 and S2). These reads were aligned to both the fox draft genome scaffolds and *de novo* brain transcriptome assembly (see Methods), producing high-quality read-count data on the 48 samples for 12,808 annotated genes in the transcriptome. Among these genes, 146 are differentially expressed in prefrontal cortex between tame and aggressive individuals at a 5% false discovery rate (*q*-value < 0.05; Figure 1B, Table S3 and Figure S5). In addition, there were 33 differentially expressed genes in basal forebrain (Table S4).

Among these hits, the two most significant genes are *DKKL1* and *PCDHGA1* (*P*-value < 10^-8^ in prefrontal cortex and *P*-value < 10^-11^ in basal forebrain; Figure 1B), and their up-regulation in tame fox was confirmed using qRT-PCR in the same RNA-seq samples (Figure S6 and Table S5; see Methods). *DKKL1* is Dickkopf-like protein 1, which has signal transducer activity and interacts with non-canonical Wnt pathway. In the mouse brain, *DKKL1* displays region specific expression, with the highest expression level in the cortical neurons of the adult cortex (7). Little is known about the function of *DKKL1* in the brain except that it bears sequence similarity to *DKK1*, an antagonist of canonical Wnt signaling implicated in a wide spectrum of physiological processes, including neurogenesis, neuronal connectivity and synapse formation. Overexpression of *DKKL1* in ventral hippocampus but not in pre-frontal cortex was associated with increased susceptibility to social defeat stress in mice (23). *PCDHGA1* is Protocadherin Gamma Subfamily A1 gene, which encodes a neural cadherin-like cell adhesion protein. Protocadherins are known to play critical roles in the establishment and function of specific cell-cell connections in the brain, such as synapse development (24) and dendrite arborization and self-avoidance in central nervous system (25, 26). *Pcdhga1* expression was down-regulated in a learned helpless rat model, suggesting its expression might affect behavior phenotypes (27). The RNA-seq experiments identified a couple hundred differentially expressed genes and they might be responsible for the behavior phenotype changes after selection.

### Expression changes occur in serotonin and glutamate receptor signaling pathways

From previous studies of pathological aggression and anxiety in humans and other animals, there is a strong prior expectation that genes involved in several neurological receptor pathways may have altered expression levels in tame foxes. Serotonin is a neurotransmitter known to play a role in feelings of well-being and happiness in humans (28). Altered expression levels of serotonin receptors have been documented in schizophrenia and bipolar disorder patients (29). Serotonin (5-HT) and serotonin metabolite (5-HIAA) levels had been found to be significantly elevated in the tame compared to the aggressive foxes (7), similar to other mammals and invertebrates (19,20). In this study, we examined genes in the serotonin receptor pathways based on the KEGG database (30, 31) and found significantly differentially expressed genes, including serotonin receptors 5A, 3A and 7, and a pair of downstream signaling genes: DUSP1 in the cAMP/PKA pathway and AKT1 in the PI3K/AKT pathway (Figure 2A and Figure S7). Nearly all the changes are in the direction of increased serotonin signaling in the tame animals.

**Figure 2.**
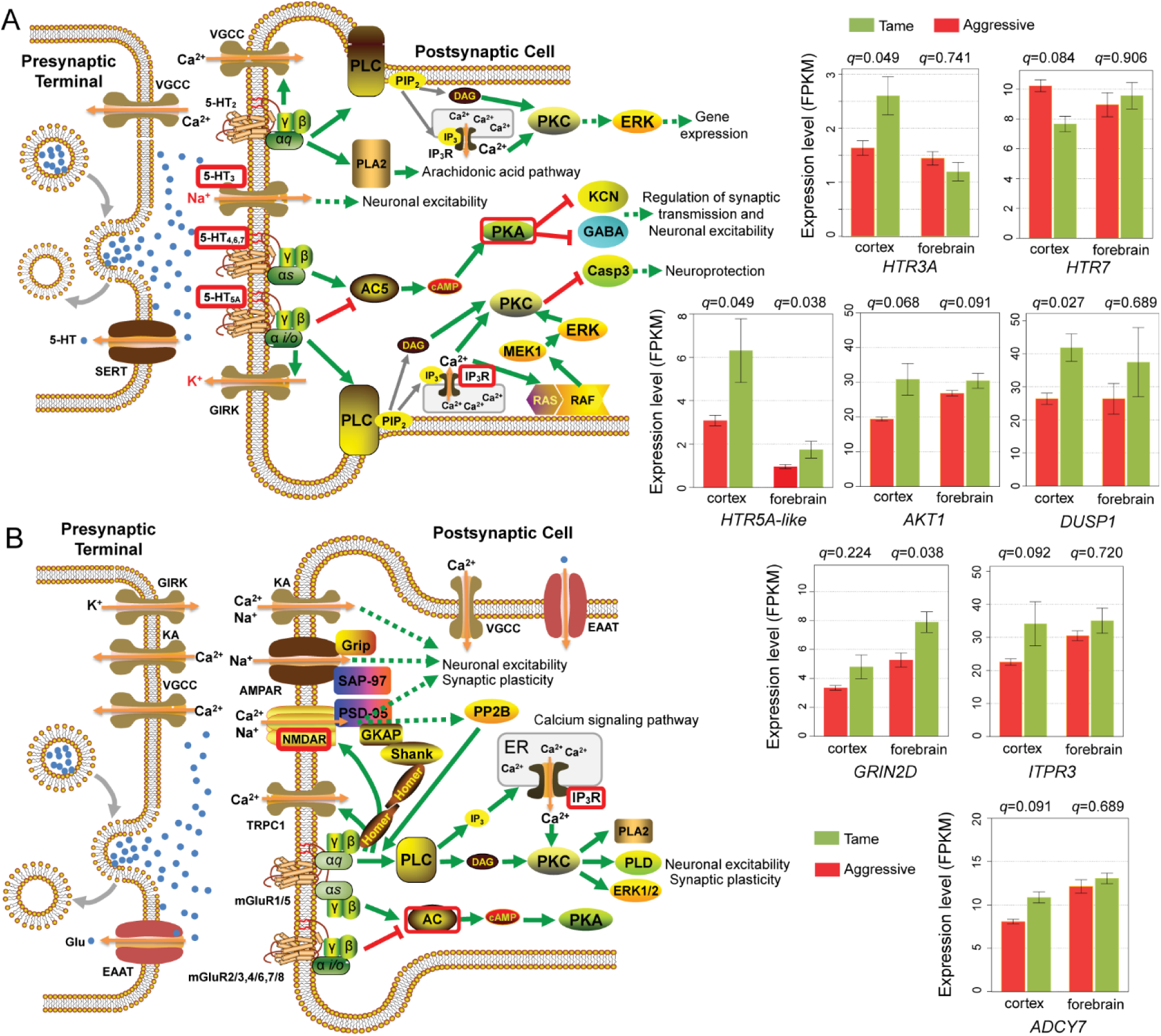
Genes that are differentially expressed between tame and aggressive fox populations in serotonin and glutamate receptor pathways. Diagrams of a serotonergic (**A**) and a glutamatergic (**B**) synapse showing the presynaptic and postsynaptic terminals (adapted from KEGG pathway database). The RNA-seq expression levels in both tissues are plotted in individual barplots for significantly differentially expressed genes (*q*-value < 0.10 in at least one tissue) between tame and aggressive foxes. Differentially expressed receptors and genes involved in downstream signaling pathways (assigned by KEGG, fig. S7 are labeled with red boxes. (**A**) In tame individuals, serotonin receptors *HTR5A-like* is up-regulated in both tissues. *HTR3A* is upregulated only in prefrontal cortex and *HTR7* is down-regulated in cortex. *DUSP1* is in cAMP/PKA pathway (labeled with red box in the middle right part of the figure) and *AKT1* is a major component of the PI3K/AKT pathway (labeled with red box in the bottom right of the figure). They are both up-regulated in tame foxes. (B) A subclass of glutamate receptors, NMDA receptor 2D (*GRIN2D:* glutamate receptor, ionotropic, N-methyl-D-aspartate 2D) and downstream signaling genes *ITPR3* and *ADCY7* (pathways labeled with red boxes in the middle right and bottom right part of the figure respectively), are differentially expressed between tame and aggressive foxes, with up-regulation in the tame animals.

Besides the critical role of serotonin, dopamine and glutamate were also known to be linked with aggression (32). In our dataset, no genes in the dopamine receptor pathway were identified to be significantly differentially expressed. For the glutamate receptor pathway, NMDA receptor 2D subunit and downstream signaling genes ITPR3 and ADCY7 were significantly up-regulated in the tame animals (Figure 2B and Figure S7). N-methyl-D-aspartate (NMDA) receptors are a subclass of glutamate receptors important for synaptic plasticity, learning and memory. This pathway also plays a key role in fear conditioning (33). Up-regulation of NMDA signaling might be consistent with increased responsiveness to keepers in the tame foxes. These results suggest that the gene expression response to selection for tameness in silver foxes impacts neurotransmitter receptor pathways, and the data sheds light on the biological basis of affiliative and aggressive behaviors by relating to neurological and pharmacological correlates with those behaviors.

### Allele frequency changes during the selection process for tame and aggressive behavior

In addition to the expression response, other genes may manifest changes in coding sequences that could affect protein function. Such genes often show allele frequency changes in their coding SNPs. In the RNA-seq data, we identified 31,025 high quality exonic SNPs (see Methods) and tested allele frequency differences at these positions.

Founder effect, inbreeding and random genetic drift can all result in allele frequency changes, and these factors need to be controlled to accurately assess the role of selection. The tame and aggressive fox populations were selected solely for specific behavioral traits, and full pedigree data for the tame (6,670 individuals) and aggressive (1,863 individuals) populations were maintained during the entire breeding program (Figures S2-S3) (11). Efforts were made to avoid close inbreeding in these populations, allowing a continuous selection for many decades and generations (9, 11). By taking advantage of this information, we directly simulated the precise effect of genetic drift and inbreeding on allele frequency changes by “gene dropping”, a method that uses the known pedigree structures for an ascertained sample of genotypes drawn from the population (in this case, the 24 RNA-seq individuals) (Figure 3A and Figure S8). At an adjusted *P*-value of 0.01, 295 SNPs in 168 genes have significantly different allele frequencies between the tame and aggressive populations (Figure 3B and Table S6), with a mean allele frequency difference of 0.79. Non-synonymous SNPs are slightly enriched in the significance of allele frequency changes compared to all exonic SNPs (25.9% vs. 23.9%), but the difference does not achieve statistical significance (Figures S9C and D).

**Figure 3.**
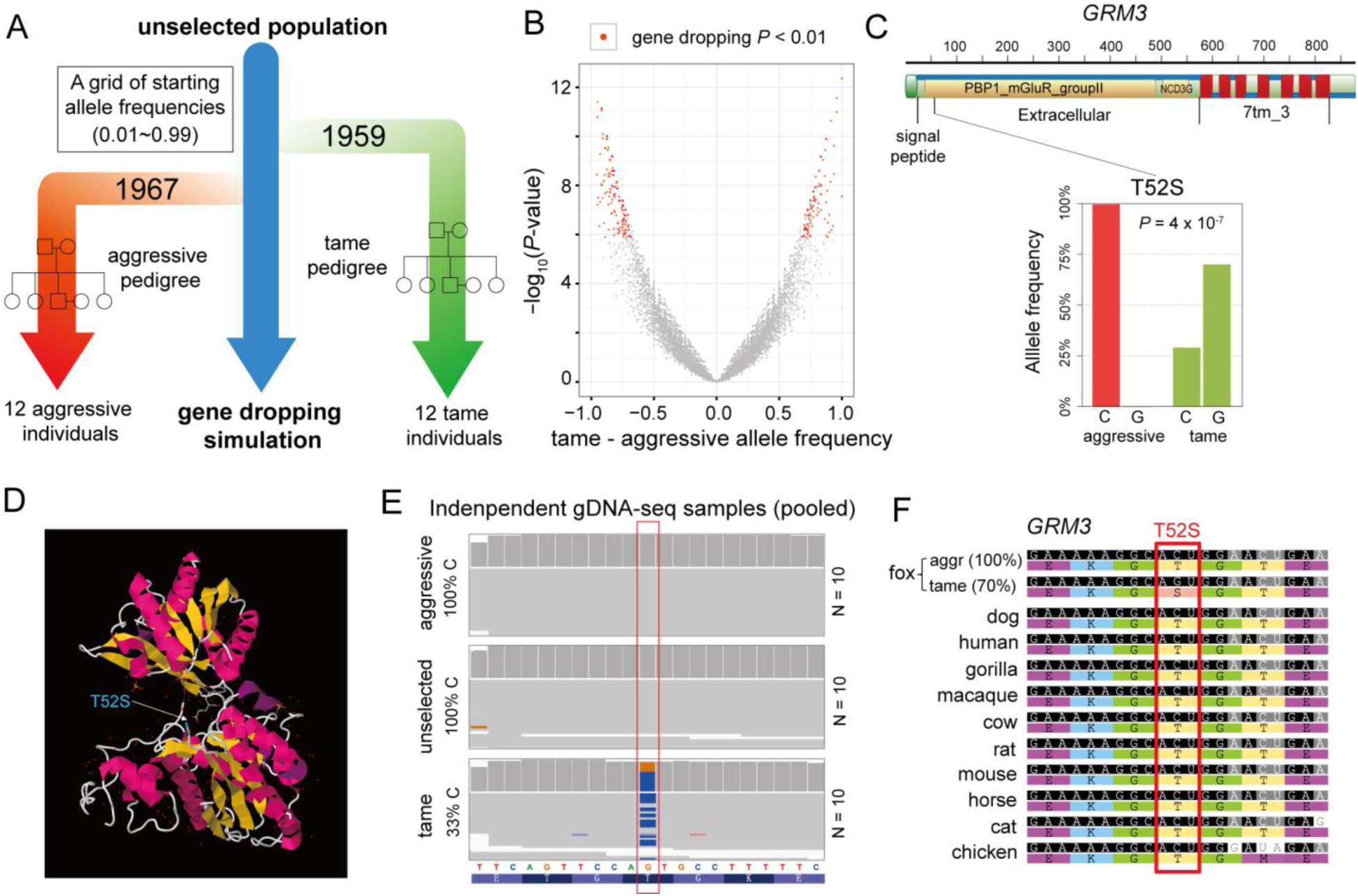
*GRM3*, a metabotropic glutamate receptor gene with significant allele frequency changes in tame population. (**A**) Gene dropping simulation scheme to determine the adjusted P-value under genetic drift, inbreeding and founder effect. A null distribution assuming no association between SNP genotypes and behavior phenotypes was generated by simulating all founder genotypes under a grid of starting founder allele frequencies (0.01~0.99 in increments of 0.01). Then alleles were dropped down the observed tame and aggressive pedigree structures (figs. S2 and S3) based on Mendelian inheritance. This was repeated many times to produce a null distribution of the magnitude of allele frequency changes. From this we obtained *P*-values for the observed allele-frequency difference between tame and aggressive RNA-seq samples. 295 SNPs are significant across all starting allele frequencies at a 1% level based on 10,000 simulations. (**B**) A volcano plot showing allele frequency difference between tame and aggressive RNA-seq samples on the *x*-axis and the –log_10_ *P*-value on the *y*-axis. Significant SNPs are labeled in red. (**C**) *GRM3* (metabotropic glutamate receptor 3) has a C→G non-synonymous SNP change causing a Thr to Ser missense mutation (T52S). In the RNA-seq data, aggressive foxes have 100% C allele and tame foxes only have 30% C allele (*P*-value = 4 x 10^-7^ and adjusted *P*-value < 0.01). PBP1_mGluR_groupII: ligand binding domain of the group II metabotropic glutamate receptor; NCD3G: Nine Cysteines Domain of family 3 GPCR; 7tm_3: 7 transmembrane sweet-taste receptor of 3 GCPR. Annotation from RCSB PDB protein data bank (ID: Q14832). (**D**) Crystal Structure of the *GRM3* extracellular region (RCSB ID: 3MS9) viewed by jmol software. T52S (labeled in blue) is near the ligand binding site, suggesting it might alter the protein function. (**E**) IGV screen shot at the *GRM3* SNP position in pooled gDNA-seq samples (figs. S10 and S11). In independently selected gDNA resequencing samples, the tame G allele frequency (67%, - strand shown in this plot) is confirmed in the tame population, and it is missing in the aggressive population. (**F**) The C allele is conserved in dogs, other mammals and the chicken. The tame G allele is the derived allele.

Ten whole-genome sequences were obtained for each of the tame, aggressive (Figures. S10-S11) and conventional farm-bred fox populations (34) at 25x coverage per population, allowing independent cross-validation of allele frequency changes. Overall, the SNP allele-frequency changes were significantly correlated (Spearman correlation coefficient ρ = 0.73, *q*-value < 0.01) between our RNA-seq and these whole-genome sequences (Figure S9B). *SorCS1*, a transporter important for trafficking AMPA glutamate receptors to the cell surface, is one of the QTL positional candidate genes with decreased heterozygosity and increased divergence between populations which was identified in the analysis of re-sequenced genomes (34). Six *SorCS1* coding SNPs are among the 295 SNPs with significant tame vs. aggressive allele frequency difference, including the third most significant SNP in the list (Table S6), highlighting the consistency of allele frequency divergence. Despite the consistent allele frequency change that occurred in *SorCS1* due to selection, no change in expression level was detected.

One of the 168 genes having a significant SNP frequency change is *GRM3*, the metabotropic glutamate receptor 3. This glutamate receptor is shown to be associated with schizophrenia, bipolar, mood disorders and delayed sexual maturity in human studies (35, 36). In our exonic SNP data, *GRM3* has a C to G change causing a Threonine-to-Serine missense mutation (T52S) in the coding region, with 100% C in the aggressive foxes and a C frequency of only 30% in the tame foxes (*P*-value = 4 x 10^-7^ and adjusted *P*-value < 0.01; Figure 3C). The altered amino acid is in the extracellular region near the glutamate binding site, which might affect the binding affinity (Figure 3D). The allele frequencies were validated in independently selected tame, aggressive and unselected individuals (Figure 3E and Figures S10-S11). The tame allele (G) is missing in both aggressive and unselected foxes. Evolutionarily the ligand binding region is highly conserved, with all genome-sequenced mammals and chicken having the C allele (Figure 3F). The increased G allele frequency might be the direct response to the artificial selection for tameness in the farm fox experiment.

### Comparative analysis with aggressive rat selection experiments and wild cat domestication revealed hits on the same genes and gene families

Our results showed that both gene expression and allele frequency responses in the tame foxes occurred in the glutamate receptor signaling pathway (genes *GRIN2D* and *GRM3*). This same pathway also experienced significant changes in both ancient domestication events as well as in recent selection experiments in other mammals. The parallel with the domestic dog is particularly noteworthy, with genes in glutamate receptor signaling (*GRIA1*, *GRIN2A*) also showing significant changes in the course of domestication (37). Similarly, in the domestication of the cat, three glutamate receptor genes, *GRIA1* and *GRIA2* were also found to be under positive selection (38). A recent selective sweep was also found in *GRIK2* in domestic rabbits (6). This convergence of selection signals on glutamate receptor signaling strongly motivates additional experimental confirmation of a functional role for glutamate signaling in behavioral differences of domesticated mammals.

Similarly, genes in the protocadherin family also display both expression and allele frequency changes during selection for tameness in foxes. Three protocadherins, *PCDH9*, *17* and *20* all have multiple SNPs with significant allele frequency changes (adjusted *p*-value < 0.01). *PCDHGA1*, a protocadherin gamma gene, is the second most significant differentially expressed gene between tame and aggressive fox brains (Figure 1B). Remarkably, another member of the same protocadherin gamma subfamily A, *Pcdhga11*, is in the list of genes associated with tameness in the rat (39). Comparative genomic analysis between domestic and wild cats also identified protocadherin A1 and B4 (*PCDHA1* and *PCDHB4*) under the selection peaks (38), suggesting a shared role of protocadherins in tame phenotypes across multiple mammalian species.

A recent QTL and transcriptome study using an F2 population of two outbred rat lines selected for tameness and aggression identified four top contributor genes for the behavior difference (39). Two (*Gltscr2* and *Lgi4*) of the top four rat candidate genes (*Gltscr2*, *Lgi4*, *Zfp40*, and *Slc17a7*) have informative SNPs in the fox data. Two synonymous coding SNPs in *Lgi4* both showed significant allele frequency differences at an adjusted *P*-value < 0.05 (table S7). Two non-synonymous and three synonymous SNPs were found in *Gltscr2*, and they were marginally significant, with an allele frequency difference of 0.375 (adjusted *P*-value = 0.10, Table S7). In sum, selection for tame/aggressive phenotypes in different mammals can lead to expression and genetic changes in genes in the same pathways.

Charles Darwin, along with many others, observed that selection for domestication in mammals often leads to a collection of phenotypes including shortened snout, curly tail, white spotting of fur on the chest, and floppy ears, often referred to as the “domestication syndrome.” These features all seem to occur in tissues that are derived from neural crest cells, suggesting that the process of selection for domestication impacts neural crest cell function (40). Intriguingly, several of the genes that manifested significant allele frequency changes in our tame foxes may play a role in neural crest cell fate (41). Wnt-signaling plays a key role in initial neutral crest cell differentiation, and both *Wnt3* and *Wnt4* in the fox had more than one SNP with significant allele frequency changes. Protocadherins are also important in neural crest cell function. Direct assessment of whether these genes play a role in neural crest cell function in the fox presents an interesting experimental challenge. In summary, the changes in expression level and allele frequency might be the direct response to the artificial selection and will help understand the genetic basis of the mammalian domestication process.

## Methods

### Brain tissue selection and dissection

Brain tissue samples were collected from adult foxes maintained at the experimental farm of the Institute of Cytology and Genetics (ICG) in Novosibirsk, Russia. All animal procedures at the ICG complied with standards for humane care and use of laboratory animals by foreign institutions. The study was approved by the Institutional Animal Care and Use Committees (IACUC) of Cornell University and the University of Illinois at Urbana-Champaign. Samples were collected from 12 foxes from the tame population and 12 foxes from the aggressive population (Figures S2 and S3). All foxes were sexually naive 1.5-year old males which were born in March to early April of 2009 and raised in the standard conditions (42). The samples were collected in August of 2010. Foxes were euthanized using sodium thiopental and brain samples were dissected immediately thereafter. The brains were cut in the sagittal plane into right and left halves and all samples were dissected from the right half. Samples from two brain regions were used in the current study: (i) prefrontal cortex; (ii) the rostral part of the basal forebrain. All samples were collected by the same scientist in a standard manner. The dissected brain samples were immediately placed into containers with RNAlater (Qiagen, Valencia, CA) and stored at −80 C.

### RNA-seq experiments and expression analysis

Total RNA samples were extracted from all 48 brain samples with Qiagen RNeasy Lipid Tissue Mini Kit (Qiagen, CA). QIAzol Lysis Reagent was used to remove excessive lipids in the brain tissue. A260/A280 absorption ratios and RNA concentrations were measured with a NanoDrop ND-1000 Spectrophotometer (Thermo Scientific, DE). RNA- seq libraries were constructed from 1.5 μg total RNA using Illumina TruSeq RNA Sample Preparation Kits v2 according to the manufacturer’s protocols (Illumina Inc., CA), and sequenced on an Illumina HiSeq2000 instrument. Single-end 50 bp reads were generated. The RNA-seq data were deposited in GEO under accession no. GSE76517.

The RNA-seq reads were aligned to both fox genome scaffolds and the transcriptome contigs from *de novo* assembly. On average 3.51% of reads had low quality or contained adapter sequence and were filtered out using Trimmomatic software (43). The fox draft genome assembly contains 676,878 scaffolds, and ones that are less than 150 bp in length and with fewer than 5 RNA-seq reads mapped were excluded from the analysis. The RNA-seq reads were mapped to the 12,851 leftover genome scaffolds using TopHat v2.0 (44). On average, 97.6% of the reads were mapped to the fox genome scaffolds and 86.9% were mapped uniquely. Two samples (488 and 490) with significantly lower mapping rate were excluded from the expression analysis. Read counts mapped to each gene model were summarized by Cufflinks v2.1.0 (45).

To get the fox transcript models and potential alternative splicing variants, we also performed *de novo* assembly of the fox brain transcripts with 1.8 billion RNA-seq reads using Trinity (46). rRNA and mtDNA reads were filtered out by custom scripts before assembly. Among the 321,151 assembled transcripts, short repetitive contigs due to gene families and repetitive sequences were removed by repeat masking and BLAT within them. The transcripts were then annotated by blasting against dog Ensembl transcripts.We compared the transcript length with 454 fox brain transcript assembly (47) and 90% of the time the Illumina assembly was longer. Among 15,551 annotated transcripts, 7,975 covered more than 80% of the orthologous dog Ensembl transcript in length, suggesting most brain transcripts were assembled close to full length. The RNA-seq reads were mapped to the transcript contig sets by BWA (48) with a maximum of 4 mismatches. Uniquely mapped read counts were summarized on annotated transcripts using BEDTools (49, 50). Genes that were differentially expressed between tame and aggressive individuals in the two brain tissues were detected with the edgeR package in Bioconductor (51, 52) at a 5% FDR level (false discovery rate, *q*-value<0.05). Normalization and expression level estimation (FPKM: <u>F</u>ragments <u>P</u>er <u>K</u>ilobase-pair of exon <u>M</u>odel) were also calculated using edgeR.

### qRT-PCR validation of selected differentially expressed genes

To confirm the RNA-seq calls of differentially expressed genes between tame and aggressive foxes, we performed qRT-PCR experiment on selected candidate genes in all 48 individual samples with two independent technical replicates (Figure S6). The tested genes were selected from the top candidate list (*PCDHGA1* and *DKKL1*) and the significant genes involved in serotonin and glutamate receptor pathways (*DUSP1*, *HTR5A-like*, *AKT1*, *ITPR3*, *GRIN2D* and *ADCY7*). qPCR primers were designed across different exons and not to overlap SNP positions between tame and aggressive populations to minimize amplification bias (Table S5). cDNAs were synthesized using SuperScript III Reverse Transcriptase (Life Technologies, CA). 10 or 100 ng total RNA were used per 15 μL qPCR reaction, depending on the signal for each gene. qPCR reactions were performed on a Roche LightCycler 480 Real-Time PCR System (Roche Diagnostics, Germany) with SYBR Green (Invitrogen, Cat No. S7563) in 384-well plates. Initial analysis was done using the Roche LightCycler 480 Relative Quantification Software. Three house-keeping genes without expression difference between tame and aggressive populations (*TBP*, *RPL14* and *EIF3D*) were selected as positive controls, and we built a standard curve using a dilution series with 4-fold increments and a total of eight data points, with two technical replicates for each point. Relative quantification was performed based on the standard curve.

### SNP calling and allele frequency estimation from RNA-seq data

To detect allele frequency changes after selection for tame and aggressive populations, we called 100,348 exonic SNPs *de novo* from the combined RNA-seq alignments on positions with 100X or more coverage depth using SAMtools (53). Local realignment over indel positions were done using GATK (54). After stringent quality filtering with custom scripts, SNP calling was performed on all 48 individual samples at 77,153 high quality SNP positions. SNPs with missing data in 7 or more individuals in tame or aggressive population were excluded, and only concordant SNPs calls in both tissues for the same individual were included in the final analysis. We also applied a cut-off restricting the analysis in SNPs with 10X or more read depth in each individual sample. For the 31,025 leftover SNPs, allele frequencies were estimated by the proportion of references alleles in each population (Figure S9).

### Pedigree analysis and gene dropping simulations

The entire tame and aggressive pedigrees were constructed based on individual data record cards from the fox farm (9, 11). The tame pedigree (offspring born year ranging from 1959 to 2010) contains 6,670 individuals including 198 founders (Figure S2). The aggressive pedigree (offspring born between 1967 and 2010) contains 1,863 individuals including 143 initial founders (Figure S3). Inbreeding coefficients for RNA-seq and gDNA-seq samples were calculated using Pedigree Viewer 6.5 (55). The entire tame and aggressive pedigrees were plotted using the PEDANTICS package in R (56). The presence of potential second sire is excluded from the analysis, because the proportion is small: 6 (0.32%) in the aggressive pedigree and 144 (2.15%) in the tame pedigree, and none of these individuals had substantial genetic contribution to the 24 RNA-seq samples. To determine the statistical significance of the allele frequency differences between tame and aggressive populations, Fisher’s exact test was used to calculate the nominal *P*-values at exonic SNP positions. Since genetic drift and a founder effect can affect allele frequencies in the pedigree, we assessed the adjusted *P*-values by directly simulating the precise effect of these confounding factors on allele frequency changes using gene dropping (57, 58). To generate a null distribution of allele frequency differences estimated from the tame and aggressive individuals under the assumption that the allele frequency dynamics are entirely determined by random drift (and hence that the SNP locus is not associated with the behavior phenotype), we first simulated all founder genotypes for tame and aggressive pedigrees according to a grid of initial allele frequencies in the conventional population (from 0.01 to 0.99 with an increment of 0.01). Then the genes were “dropped” down both pedigrees based on Mendelian inheritance and performing a random draw for gametes transmitted by heterozygotes. Allele frequencies were calculated for the 12 tame and 12 aggressive RNA-seq samples, and the test statistic is the allele-frequency difference. We simulated this entire process 10,000 times to obtain null distributions for the test statistic under all possible initial founder allele frequencies (Figure. S8). The SNP is significant at a 1% level if the observed allele frequency changes were greater than all the expected ones under the null hypothesis for all possibl starting allele frequencies.

## Acknowledgements

We gratefully acknowledge the Meinig Family Investigator Award to A.G.C. for suppoi We thank Amanda Manfredo and Li (Grace) Chi for assistance with the experiments. W are grateful to Irina V. Pivovarova, Tatyana I. Semenova, and all the animal keepers at the ICG experimental farm for research assistance. The project was supported by National Institutes of Health grant GM120782, USDA Federal Hatch Project 538922, FCP grant 84-74, Russian Science Foundation, and FANO budget project VI.53.2.4. Th fox brain RNA-seq data were deposited in GEO under accession no. GSE76517.

